# Using an agent-based sexual-network model to guide mitigation efforts for controlling chlamydia

**DOI:** 10.1101/233239

**Authors:** Asma Azizi, Jeremy Dewar, James M. Hyman

## Abstract

We create and analyze a stochastic heterosexual agent-based bipartite network model to help understand the spread of chlamydia trachomatis. Chlamydia is the most common sexually transmitted infection in the United States and is major cause of infertility, pelvic inflammatory disease, and ectopic pregnancy among women. We use an agent-based network model to capture the complex heterogeneous assortative sexual mixing network of men and women. Both long-term and casual partnerships are modeled with different sexual contact frequencies and condom use. We use simulations to compare the effectiveness of intervention strategies based on randomly screening people for infection, treating the partners of infected people, and rescreening for infection after treatment. We compare the difference between treating the partners of an infected person both with, and without, testing them first for infection. The highest prevalence is among young sexually active individuals. We calibrate the model parameters to agree with recent survey data showing chlamydia prevalence of 14% of the women and 9% of the men in the 15 – 25 year-old African American residents of New Orleans, Louisiana. We observed that although increased chlamydia screening and treating most of the partners of infected people will reduce the incidence, these mitigations alone are not sufficient to control the epidemic. The model predicts that the current epidemic can brought under control once over half of the partners of infected people are tested and treated.

## Introduction

Chlamydia trachomatis (Ct) is the most commonly reported sexually transmitted infection (STI) in the United States, with over 1.8 million cases each year [1]. It is a major cause of infertility, pelvic inflammatory disease (PID), and ectopic pregnancy among women [2–8], and has been associated with increased HIV acquisition [2,4,6–8]. Untreated, an estimated 16% of women with Ct will develop PID [2], and 6% will have tubal infertility [7]. In southern U.S cities, including New Orleans, there is an ongoing epidemic of Ct in young African American (AA) adults. A pilot study in this community [9] found an average of 1.5 sexual partners per person per three months, a relatively high turnover in sexual partners, and approximately 11% prevalence of Ct infection. The high prevalence along with high turnover stresses the need for more effective mitigation efforts to bring the epidemic under control.

Further complicating the issue, about 70% — 95% of women and 90% of men infected with Ct are asymptomatic and still transmit the infection to others. When Ct prevalence is high, regular screening is an effective approach to identify and treat infected individuals. Since 2000, women have been typically screened for Ct as part of their physical exam. However, men are not routinely screened for infection, therefore, untreated men may serve as a reservoir, reinfecting women after they are treated. We will investigate the impact that increased Ct screening of men in high prevalence areas on the disease prevalence. Typically, when someone is found to be infected, they are encouraged to have their sexual partners notified and tested for infection. Sometimes the partners are treated, without first being tested for infection. If a partner is also tested for infection and found to be infected, then their partners can be notified and treated, identifying a chain of high-risk individuals who might be spreading Ct.

Transmission-based mathematical models can help the public health community to understand and to anticipate the spread of diseases in different populations and to evaluate the potential effectiveness of approaches for bringing the epidemic under the control [10]. These models create frameworks that capture the underlying Ct epidemiology and the heterosexual social structure underlying the transmission dynamics. An ordinary differential equation Ct transmission model developed by Althaus et al. [11] captures the most essential transitions through an infection with Ct to assess the impact of Ct infection screening programs. Using sensitivity analysis they identified the time to recovery from infection, and the duration of the asymptomatic period, as the two of most important model parameters governing the disease prevalence.

Clarke et al. [12] investigated how control plans can affect observable quantities and demonstrated that partner positivity (the probability that the partner of an infected person is infected) is insensitive to changes in screening coverage or contact tracing efficiency. They also evaluated the cost-effectiveness of increasing contact tracing versus screening and concluded that random screening coupled with partner notification is the a cost effective mitigation approach.

A selective sexual mixing hybrid differential/integral equation Ct model was developed by Azizi et al. [13] to capture the heterogeneous mixing among people with different numbers of partners. The model stratified the population based on number of partners and used sensitivity analysis to identify the importance of mixing on the effectiveness of various mitigations, such as condom use. An alternative, and more realistic model, is to represent the sexual network by a graph where each individual within a population is a node. The connecting edges between the nodes denote sexual relationships that could lead to the transmission of infection. These sexual mixing networks can capture the heterogeneity of whom an infected person can, or cannot, infect.

Kretzschmar et al. [14] evaluated different screening and partner referral methodologies on a network in controlling Ct. They designed an stochastic model based on a stochastic pair formation and separation process. The process modeled the spread of STIs particularly gonorrhea and chlamydia in a general age-structured heterosexual population in the Netherlands and implemented several intervention scenarios including contact tracing, screening and condom-use. Their results show contact tracing and increased condom-use were more effective than screening to control the epidemic of these STIs. They observed that screening is most effective when it targets the core group of highly sexually active people.

We create an agent-based heterosexual network model of Ct transmission to evaluate potential intervention strategies for reducing the Ct prevalence in urban cities, such as New Orleans. We construct a network model that mimics the heterosexual behavior obtained from a sexual behavior survey of the young adult AA population in New Orleans and model Ct transmission as a discrete time Monte Carlo stochastic event on this network. The model is initialized to agree with the current New Orleans Ct prevalence. We use sensitivity analysis to quantify the effectiveness of different prevention and intervention scenarios, including screening, notification of partners, which includes partner treatment and partner screening (contact tracing), condom-use, and rescreening [15].

## Materials and Methods

In our heterosexual network model, each person is a distinct identity represented by a node in the network. This model structure allows the sexual partnership dynamics, such as partner concurrency, sexual histories of each person, and complex sexual networks, to be governed at the individual level [16].

The data used for our network was previously collected by two research studies in New Orleans, LA. One, a pilot study of community based STI testing and treatment for African American men ages 15 – 25 [9] and the other, a study of an Internet based unintended pregnancy prevention intervention for African American women ages 18 – 19 [17]. These two studies were reviewed and approved by the Tulane University Institutional Review Board. The 202 men and 414 women enrolled in these studies were asked for the number of different heterosexual partners they have had in the past three months (for women) or two months (for men). Women were also asked to estimate the number of partners that their partners have had in the same time period.

The survey results were used to construct the bipartite heterosexual network of 2000 men and 3000 women. The generated network statistically agrees with the distribution for the number of partners men and women have had in the past three months, and the distribution for the number of partners of their partners (the joint degree distribution) [18].

Each node **i** in the network represents a person, denoted by the index **i**, and each edge e_**ij**_ represents sexual partnership between two nodes **i** and **j**. The network is weighted, where the weight 0 < *w*_**ij**_ ≤ 1 for edge e_**ij**_ is the probability that there will be a sexual act between two partners **i** and **j** on an average day. In the model, each day and through a stochastic process, the edge **e_ij_** will exist (turn on) with probability *w*_**ij**_, the probability that two nodes **i** and **j** have sexual act in that day, or not exist (turn off) with probability of 1 – *w*_**ij**_, which is equivalent to not having an edge (sexual act) between individuals **i** and **j** on that day.

In our stochastic Susceptible–Infectious–Susceptible (SIS) model, a person **i** is either infected with Ct, *I*_**i**_(*t*), or susceptible to being infected, *S*_**i**_(*t*). During the day *t*, an infected person, *I*_**j**_(*t*), can infect any of the susceptible partners, *S*_**i**_(*t*), they have sexual contact with. We define λ_*ij*_ as the probability that *S*_**i**_(*t*) will be infected by *I*_**j**_(*t*) by the end of the day, 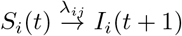. Similarly, we define *γ_j_* as the probability that an infected person, *I*_**j**_(*t*), will recover by the end of the day, 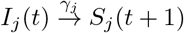.

### Force of infection

The force of infection, λ*_ij_*(*t*), is the probability that a susceptible person *S_j_* is infected on day *t* by *I_j_*. This depends on probability of a sexual act between person *i* and *j* on an typical day, as defined by edge weight, *w_ij_*, in the model. We define *β_nc_* as the probability of transmission per act when a condom is not used, and *β_c_* as the reduced probability of transmission per act when a condom is used. The forces of infection between **i** and **j** for when condom is not used, 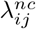, and for when condom is used, 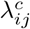, are defined by

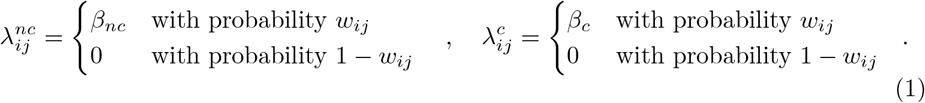

We assume that couples use a condom in *κ* fraction of their acts correctly and that the condom is 90% effective in preventing the infection from being transmitted, that is, *β_c_* = 0.1*β_nc_*.

### Recovery from infection

The model accounts for infected people recovering through natural recovery or after being treated with antibiotics. We assume that a fraction of the people treated for infection will return later to be retested for infection.

#### Natural recovery

We assume that all infected people will eventually recover and return to susceptible status, even if they are not treated for infection. In the model, the time for natural (untreated) recovery has an exponential distribution with an average time of infection of 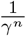 days, and the duration of infection for an individual is a random number from this distribution.

#### Recovery through treatment

We assume that the time for infected person to recover after treatment is a log-normal distribution with an average of 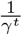 days. That is, the duration of infection for a treated infected person **k** would be set to a random number that follows log-normal distribution, 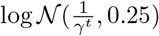, rounded to the nearest day. In the model, if that number of days is smaller than the duration remaining for naturally clearing the disease, then the shorter time is use for the recovery period.

Each year, a fraction of the population is tested for Ct infection, e.g. through a routine medical exam (random screening) or after being notified that one of their previous partners was infected.

#### Random Screening

We define random screening as testing for infection when there are no compelling reasons to suspect a person is infected. For example, random screening might be part of a routine physical exam and is an effective mitigation policy to identify asymptomatic infections. In our model, we assume that the fraction *σ_y_*% of people are randomly screened each year, and an individual is screened with probability 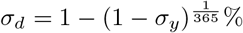 each day.

#### Notification of Partners

We assume that an infected person notifies *θ_n_* fraction of their partners about their exposure to infection. Some of the notified partners will do nothing, some will test for Ct infection, and some will seek treatment for Ct without first testing because it is simpler. The notified partners of a treated person are divided into three classes:

1. Partner Treatment: *θ_t_* fraction of the notified partners are treated, without first testing for infection.
2. Partner Screening: *θ_s_* fraction of the notified partners engage in screening test and then start treatment if infected.
3. Do nothing: (1 – *θ_t_* – *θ_s_*) fraction are not tested or treated.

For simplicity, we assume notified partners are the ones who are notified and do something, that is, we assume 1 – *θ_t_* – *θ_s_* = 0. Then we define *θ_n_θ_t_* as fraction of partners of an infected person who are treated without first testing for infection (Partner Treatment), and *θ_n_θ_s_* as fraction of the partners follow screening test (Partner Screening). The model includes a time-lag of *τ_N_* days between the day a person is found to be infected and the day their partners are notified.

Notification of partners spreads across the same network that originally spread the disease. The partner screening approach is more effective since every time the partner of an infected person is found to be infected, then the cycle repeats itself and their partners are notified.

#### Rescreening

A common practice in disease control is rescreening. People found to be infected are given a treatment and asked to return after a short period to be tested again. In our model, we assume that a fraction, *σ_r_*, of the treated people return for retesting *τ_R_* days after treatment.

### Model initialization

Our goal is to model the current Ct epidemic in New Orleans with an initial prevalence of ¿o. Infected people are not dropped into an otherwise susceptible population, instead they are distributed as they would be as part of an emerging epidemic, one that started some time in the past. We call these initial conditions *balanced* because when the simulation starts the infected and susceptible populations, along with durations of infection, are in balance as an emerging epidemic would on average have. When the initial conditions are not balanced, then there is usually a rapid (nonphysical) initial transient of infections that quickly dies out as the infected and susceptible populations relax to a realistic infection network.

To define the balanced initial conditions, we start an epidemic in the past by randomly infecting a few high degree individuals. We then advance the simulation until the epidemic grows to the prevalence *i*_0_. We then reset the time clock to zero and use this distribution of infected people, complete with their current infection timetable, as our initial conditions. Because these are stochastic simulations, when doing an ensemble of runs we reinitialize each simulation by seeding different initial infected individuals. Fig 1 illustrates the typical progression of the epidemic to reach the current Ct prevalence of 9% in men and 14% women in the 15 – 25 year-old New Orleans AA community. The numerical simulations comparing the different mitigation strategies all start at this endemic stochastic equilibrium.

**Fig 1.**
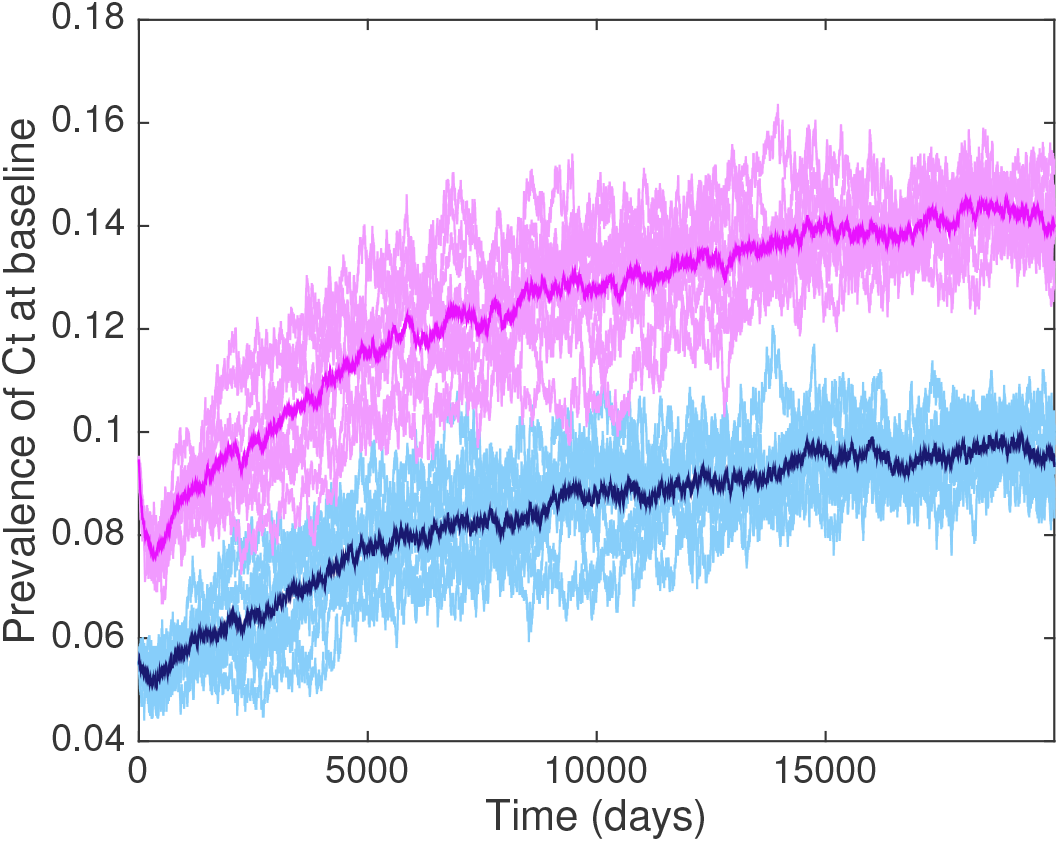
Prevalence increases to reach the current steady state. About 9% of men (blue lower curve) and 14% of women (pink upper curve) are infected at the equilibrium for the baseline model parameters. This is approximately the current prevalence in New Orleans 15–25 year-old AA population. The light areas around the dark mean values show the range of the solutions after 10 simulations.

## Results

We compare the model-projected impact of increased random screening, notification of partners – which includes partner screening and partner treatment – and rescreening on the prevalence of Ct infection. All of the simulations start at a balanced equilibrium obtained with the model baseline parameters in Table 1.

**Table 1.** **Parameters definition and value**: the model parameters describing the transmission of Ct infection, as well as recovery associated with natural recovery, screening, and notification of partners were obtained from the literature [9,11,19,20], but other parameters are calibrated to biological, behavioral, and epidemiological data from general heterosexual population resides in New Orleans. Probability of transmission per act is calibrated to a baseline prevalence of 13.95% among women and 9% among men.

| Parameter | Description | Baseline | Unit | Reference |
| --- | --- | --- | --- | --- |
| $\Delta t$ | Time step | 1 | day | – |
| $P^m(P^w)$ | Population of men (women) | 2000(3000) | people | Assumed |
| $i_0$ | Balanced initial prevalence | 0.08 | – | Assumed |
| $\beta^{m2w}$ | Probability of transmission per act from men to women | 0.16 | – | Calculated |
| $\beta^{w2m}$ | Probability of transmission per act from women to men | 0.16 | – | Calculated |
| $\kappa$ | Fraction of times that condoms are properly used during sex | 0.6 | – | [9] |
| $1/\gamma^n$ | Average time to recover without treatment | 365 | days | [11, 19] |
| $1/\gamma^t$ | Average time to recover with treatment | 7 | days | [20] |
| $\sigma_y^m$ | Fraction of men randomly screened per year | 0.05 | – | [9] |
| $\sigma_y^w$ | Fraction of women randomly screened per year | 0.45 | – | [9] |
| $\sigma_r$ | Fraction of infected people return for rescreening | 0.10 | – | Assumed |
| $\theta_n$ | Fraction of the partners of an infected person who are notified and do test or treated for infection | 0.26 | – | [9] |
| $\theta_t$ | Fraction of notified partners of an infected person who are treated without testing | 0.75 | – | [9] |
| $\theta_s$ | Fraction of notified partners of an infected person who are tested and treated for infection | 0.25 | – | [9] |
| $\tau_N$ | Time lag of partner notification | 5 | days | Assumed |
| $\tau_R$ | Time lag of re-screening | 100 | days | Calculated |

### Random screening

To determine the effectiveness of increasing the number of men screened for Ct per year, we compare the steady state prevalence by varying the fraction of men who are screened randomly each year, 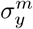. The current screening rate for young men for Ct in high prevalence areas, like New Orleans, is low. This scenario can estimate the cost effectiveness of increased screening of young men on the Ct prevalence in women [21,22]. Fig 2 shows a reduction in the overall Ct prevalence as the number of men randomly screened for Ct increases from 0 to 50%, 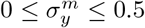. The filled circles are the mean of ten different stochastic simulations (open circles). The least-square linear fit suggests that the steady-state Ct prevalence will decrease by 0.01 for every additional 10% of the men screened during a year. Though a drop of five percent in prevalence is an admirable decrease, increased screening alone would not be sufficient to control Ct.

**Fig 2.**
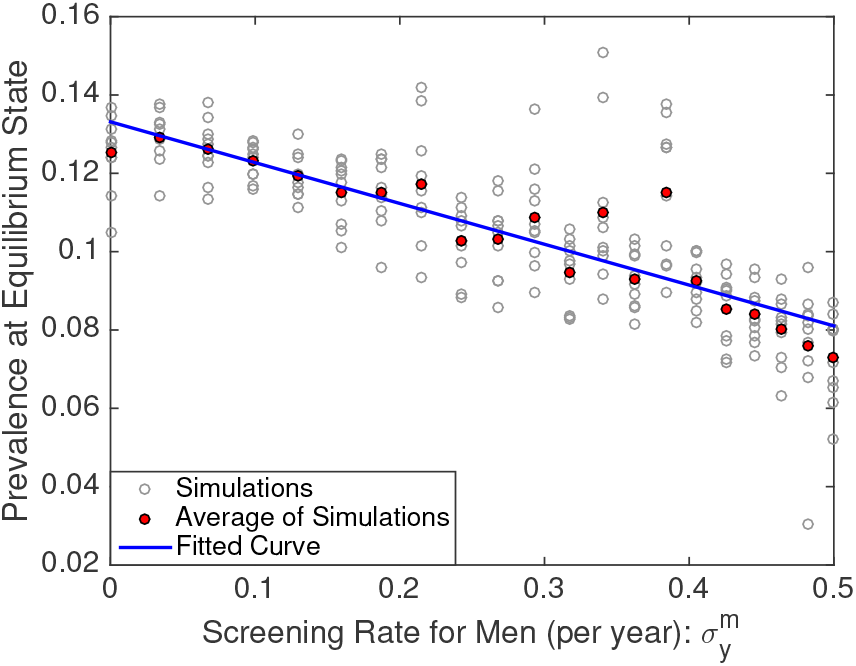
Steady state prevalence decreases as more men are screened each year. 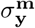: a low negative correlation, screening men randomly by 50% reduces prevalence by 5% which is not effective enough to implement as a sole intervention.

### Notification of partners

This scenario of notifying partners quantifies the impact of giving an infected person’s partners a chance to be tested and treated. We define *θ_n_* as the fraction of infected person’s partners who are notified that they might be infected. We then assume that only *θ_t_* fraction of those notified partners are treated, without testing (partner treatment), and *θ_s_* fraction are tested and if necessary, treated (partner screening). Note that the fraction 1 – *θ_n_* fraction of the partners are not notified.

#### Partner treatment

In partner treatment we assume when someone is found to be infected the fraction θn of their partners are notified and then all of the notified partners will seek treatment without testing, in other words, we define *θ_t_* = 1. Fig 3, shows the impact of partner treatment ranging from no notified partners treated, *θ_n_* = 0, to all notified partners treated, *θ_n_* = 1. The filled circles are the mean of ten different stochastic simulations (open circles). The least-square linear fit suggests that the steady-state Ct prevalence will decrease by 0.07 for every 10% increment in the fraction of notified partners seeking treatment. This practice, although common in disease control today, is not as effective as partner screening as we will see.

**Fig 3.**
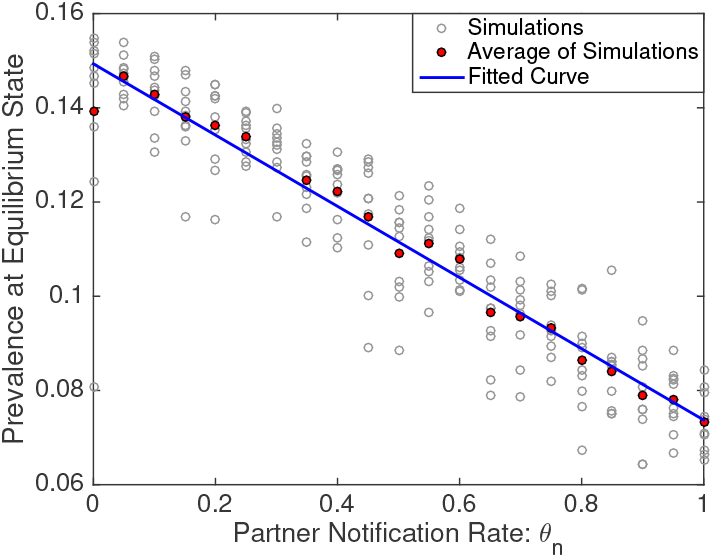
Prevalence decreases as the more partners are treated after being notified that they might be infected. In these simulations, we assume that all the notified partners are treated, without testing *θ*_**t**_ = 1. This approach is only mildly effective and the prevalence remains high (8%), even when all the partners of treated people are treated.

#### Partner screening

To quantify the impact of screening the partners of an infected person, where partners are tested then treated if found to be infected, we assume that all notified partners of an infected person are screened, that is, we assume *θ_s_* = 1. Fig 4 shows the impact as *θ_n_* varies from 0 to 1. The filled circles are the mean of ten different stochastic simulations (open circles). The least-square linear fit suggests that there is a threshold effect (tipping point) at *θ_n_* ≈ 0.4, when the approach becomes extremely effective. This happens when the partner screening percolates through the sexual network to identify the infected individuals. Our model indicates that this is by far the most effective approach for bringing the epidemic under control.

**Fig 4.**
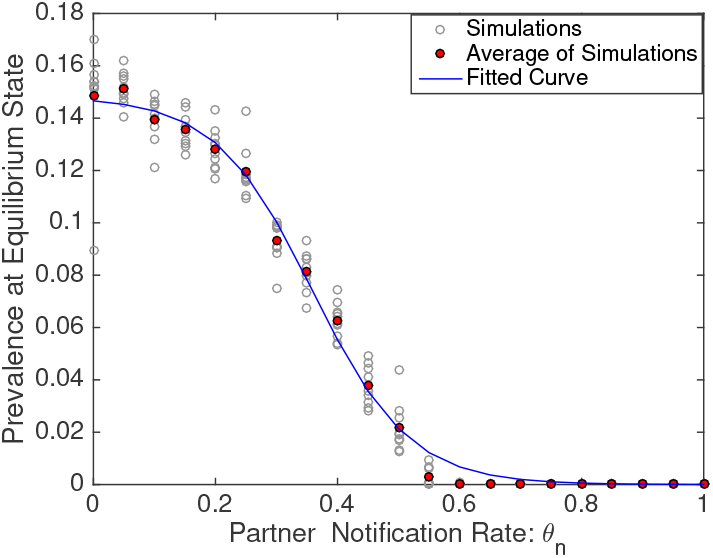
Prevalence drops to zero as the fraction of the partners of treated people are tested before possible treatment. In these simulations we assume that all of the notified partners are tested for infection, *θ_s_* = 1. This partner screening approach is highly effective if the fraction of tested partners, *θ_n_*, exceeds its critical value 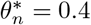. That is, when 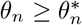 and *θ_s_* = 1, the Ct prevalence rapidly decays to zero.

#### Partner treatment and screening

In reality, some of the notified partners will seek treatment without testing, and some will allow themselves to be tested before being treated. We quantify the effectiveness of this mixture of the two previous scenarios by varying fraction of the partners taking action (*θ_n_* = 0.10,0.20, 0.50, 0.65 and 0.80), along with the fraction of these notified partners that seek just treatment (*θ_t_*) and the fraction being screened for infection (*θ_s_* = 1 – *θ_t_*).

When few partners are notified and take action (*θ_n_* is small), then partner treatment and partner screening have almost the same impact on controlling the prevalence. For example, for *θ_n_* = 0.10, the prevalence versus *θ_t_* = 1 – *θ_s_* is almost flat, that is, there is no difference between cases if partners follow treatment without testing or first test and then treat if infected.

As *θ_n_* increases the partner screening becomes a highly successful mitigation policy. Consider the case when half of the partners are notified and take action, *θ_n_* = 0.5, and half of them go for screening, *θ_s_* = 0.5, and the other half go for treatment, *θ_t_* = 0.5. That is, half of an infected person’s partners do nothing, a fraction *θ_n_θ_t_* = 0.5 × 0.5 = 0.25 are treated without testing for infection, and a fraction *θ_n_θ_s_* = 0.5 × 0.5 = 0.25 are tested and treated if found infected. If any of the tested notified partners of the infected person are found to be infected, their partners are then notified and the cycle repeats to spread out and identify more infected people. This conditional percolation of screening through the sexual network is why this policy is so effective. For this case, the prevalence reduction is 7%. Thus, compared with if all the notified partners follow treatment without testing, *θ_t_* = 1, which reduces the prevalence by only 1%, it works better. But compared with if all the notified partners follow test and treat if necessary, θ_s_ = 1, which reduces the prevalence by 11%, this combined scenario is not the one to select, Fig 5.

**Fig 5.**
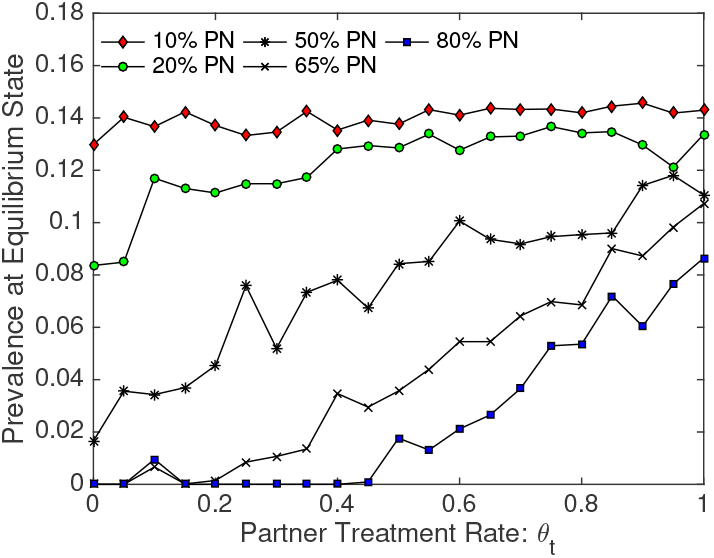
Prevalence at steady state increases when the fraction of partners notified are treated and not tested. When only a few partners of an infected person are notified, *θ_n_* is small, then partner treatment and partner screening have similar small impact on Ct prevalence. When more partners of infected people take action, *θ_n_* increases, then the partner screening strategy is more effective in controlling the infection.

It is important to note that partner screening is more expensive than partner treatment. Given this last scenario, this suggests that when the fraction of partners we are able to notify, *θ_n_*, is small then partner screening may not be a good strategy compared to partner treatment. However, if a large enough fraction of partners are notified then it is better to test and treat (partner screening) to control the spread of Ct effectively.

### Rescreening

The rescreening scenario targets two goals: first finding the time-lag for resceering, second quantifying the impact of rescreening on prevalence of Ct.

#### Interval for rescreening

People who are found to be infected are more likely to be reinfected in the future. Repeated Ct infection can be the result of treatment failure, sexual activity with a new partner, or being reinfected from an existing infected partner. It makes sense to ask the infected people who were treated to return in a few months for retesting. We will use the model to compare the rates of reinfection to help optimize the time, *τ_r_*, from treatment to rescreening.

The time *τ_r_* for rescreening should be long enough so it is likely that the person will be reinfected, but not so long that such a reinfected person could infect significantly more people. We start by identifying the rescreening time when the prevalence for the treated population exceeds the prevalence for the whole population. That is, it becomes cost effective to rescreen when over 15% of previously screened people are again infected. To find this optimal time, we compute the time taken between screening time and next reinfection time for all individuals in the network assuming there is no rescreening i.e *σ_r_* =0. On average at baseline 13% of individuals are infected, therefore, through random screening, 13% of infected individuals are found. The current CDC guidelines recommend that people are rescreened for infection three months after treatment [23].

We note that in our model, a person may receive screening test and get reinfected more than one time, therefore, we count the number of tests and reinfection events rather than the number of individuals with a test and infection.

We plot cumulative distribution of time between screening and reinfection events in Fig 6. Past studies have observed that about 25% of the rescreened people are again found to be infected by three months. The Figure demonstrates that in our model also predicts that about 25% of treated individuals are again infected after 100 days. Although the model supports the CDC guideline as reasonable, the time between treatment and rescreening could be shortened to two months with an improved impact.

**Fig 6.**
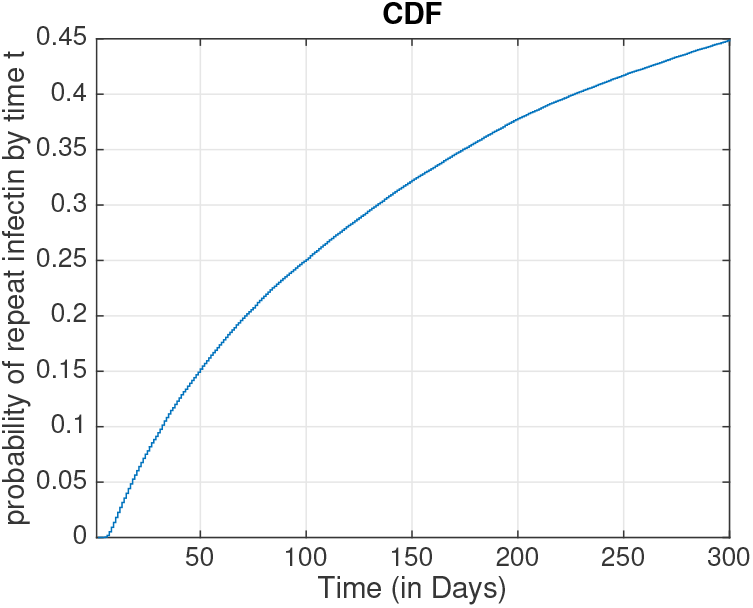
Truncated cumulative probability distribution of time between treatment and reinfection with Ct. About 25% of the treated people are again infected after 100 days. This increases to about 45% are reinfected after a year.

#### Rescreening rate

The secondary goal of rescreening scenario is to determine if rescreening for Ct infection in a larger rate would be successful to reduce its prevalence. At the baseline case only 10% of screened individuals return for rescreening. We assume 0 ≤ *σ_r_* ≤ 1 fraction of screened individuals participate in rescreening plan 100 days after their current screening day. The Fig 7 quantifies the prevalence of Ct at steady state dependent on rescreening rate *σ_r_*: there is a negative correlation between prevalence at steady state and *σ_r_* when *σ_r_* fraction of screened individuals returns for rescreening, if *σ_r_* fraction of screened individuals follow screening again then the prevalence reduces roughly by 0.02 *σ_r_*. Although there is a high chance of reinfection when the individual’s behavior does not change, we do not observe an effective impact on prevalence of Ct by monitoring infected individuals. Rescreening program has a trend similar to screening, and none of them are effective as sole intervention because they are not able to find the chain of infection like partner screening. On the other hand sensitivity of prevalence to rescreening is less than that of screening, indicating the fact that for a limited budget the idea of finding more people to screen, random Screening, is more effective than frequent screening for less people.

**Fig 7.**
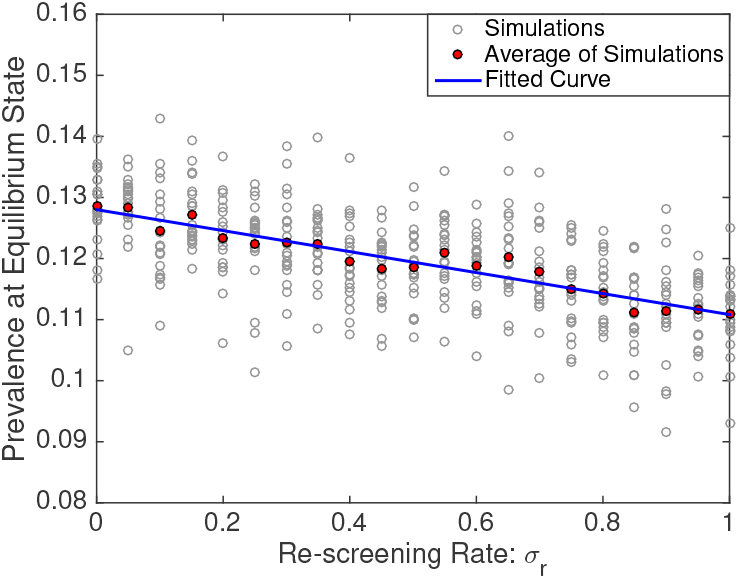
Prevalence at steady state is relatively insensitive to the rescreening rate *σ_r_*. Even rescreening all the infected people reduces prevalence by only 2%. This is a cost effective approach, but must be used on conjunction with other mitigation efforts.

## Discussion

We used heterosexual behavior survey and Ct prevalence data for the young adult AA population in New Orleans to create a stochastic, Monte Carlo - Markov Chain, agent-based bipartite sexually transmitted disease-transmission network model. In the model, men and women are represented by the network nodes and sexual partners are characterized by edges between the nodes. The edges between partners in the network dynamically appear and disappear each day depending if the individuals have sexual act on that day. The joint degree distribution of the network captures the correlation of an individual’s risk (their number of partners) with their partner’s risk (number of partners of their partners). Our network model is updated each day to account for sexual acts as a dynamic variable, which is an improvement over previous network models.

We use to model to quantify the impact of increasing screening men for infection, partner notification, and rescreening of treated individuals on reducing Ct prevalence. We observed that increasing Ct screening of men has only a modest impact on reducing Ct prevalence in the young adult AAs in New Orleans, Fig 2. Starting at a baseline of 13% prevalence under the assumption that 45% of the women are being screened each year for Ct, then increasing the screening of men from 0% to 50% would only reduce the overall Ct prevalence to 8%. That is, increasing Ct screening of men alone would not bring the epidemic under control.

In evaluating the effectiveness of notification of partners we assumed that a fraction of the partners of an infected person will seek treatment (without testing) or be screened (tested and treated) for infection. We observed that if most of the notified partners were treated, without testing, then this mitigation had only a modest impact on Ct prevalence. However, when the partners of an infected person were tested before treatment, there was a tipping point where partner screening would bring the epidemic under control. That is, when over 40% of notified partners of all the infected people are screened for infection, then the Ct prevalence rapidly decreased to very low levels Fig 5. This critical threshold represents the partner screening level where a contact tracing tree can spread through the heterosexual network to identify and treat most of the infected people.

In rescreening, an infected individual returns for testing a few months after they are treated. We used the model to estimate the probability that a treated person would be reinfected as a function of the time since they were treated. The CDC guidelines recommend that treated people return for screening three months after treatment. We observed that for the baseline case of 13% infected population, about 25% of the treated population were reinfected three months after treatment. We observed that although the rescreening is a cost effective approach to identify infected people it has only a small impact on the overall Ct prevalence.

Our bipartite heterosexual network model was constructed based on the correlations between the number of partners a person has and the number of partners their partners have. The model can be made more realistic by embedding this sexual network within a social network that includes casual relationships. The social network would capture correlations between relationships based on age, where people live, and economic status.

Also, although our model includes condom use, it does not account for behavior changes, such as increased condom use after being treated for infection or the differences in condom use between primary and casual partners.

Our future research will improve the model so we can better quantify the impact of counseling and behavioral changes such as increasing condom use or partner notification rates.

## Acknowledgments

The authors thank Patricia Kissinger and Norine Schmidt for their useful comments and suggestions. This work was supported by the endowment for the Evelyn and John G. Phillips Distinguished Chair in Mathematics at Tulane University and grants from the National Institutes of Health National Institute of Child Health and Human Development, NCHID/NIAID ( R01HD086794) and Office of Adolescent Health, OAH (TP2AH000013) and the National Institute of General Medical Sciences program for Models of Infectious Disease Agent Study (U01GM097658). The content is solely the responsibility of the authors and does not necessarily represent the official views of the National Institutes of Health.

